# When Randomness Becomes Rigid: Dynamic Connectivity Entropy and Symptom-Linked Network Dysfunction in Schizophrenia

**DOI:** 10.64898/2026.01.18.700221

**Authors:** Natalia Maksymchuk, Robyn L. Miller, Armin Iraji, Vince D. Calhoun

## Abstract

High dimensionality of dynamic functional connectivity (dFNC) data representation complicates clinical interpretation and biomarker discovery. We propose a new complementary analytical framework based on dynamic inter-network connectivity entropy (DICE) and its potential use as a biomarker of mental illness. Our framework shows that DICE features extend beyond patient–control discrimination, revealing distinct pathophysiological signatures and differential associations with symptom dimensions.

Using resting-state fMRI data from 311 participants, 160 controls, 151 schizophrenia (SZ) patients, we identified 53 intrinsic networks, computed DICE and derived three families of DICE-based metrics: (i) entropy level and range, (ii) distributional shape and temporal organization, (iii) entropy-state repertoire and occupancy. These measures revealed a multidimensional signature of altered entropy dynamics in SZ: (1) elevated baseline entropy with reduced fluctuation magnitude and reduced entropy acceleration; (2) reduced temporal persistence of entropy excursions and entropy distributions closer to Gaussian; and (3) a narrowed repertoire of entropy states, prolonged time in near-baseline entropy configurations. The DICE-based metrics within the SZ group show different associations with symptom dimensions. Reduced fluctuation magnitude and acceleration were associated with greater PANSS general symptom severity (disturbance of volition and preoccupation). Reduced deviation from Gaussianity was associated with higher PANSS positive severity (delusions and hallucinations). Reduced temporal persistence was associated with multiple PANSS positive, negative, and general symptoms. Reduced entropy-state diversity and prolonged dwell time in near-baseline states were associated with depression and PANSS positive/general severity, respectively. The multidimensional pathophysiology revealed through the different entropy patterns may potentially guide biomarker development and personalized treatments.

## Introduction

Schizophrenia (SZ) is a severe psychiatric disorder causing substantial functional disability (Charlson et al., 2018; Same et al., 2024) and economic burden (Calzavara Pinton et al., 2024; Kadakia et al., 2022). Dysconnectivity hypothesis proposes that SZ arises from disrupted functional integration across distributed brain networks (Bullmore et al., 1997; Calhoun et al., 2009; Calhoun et al., 2011; Stephan et al., 2009), supported by convergent evidence of reduced white matter integrity (Ellison-Wright & Bullmore, 2009), altered gray matter density (Glahn et al., 2008), and aberrant functional connectivity patterns showing both hyperconnectivity and hypoconnectivity (Calhoun et al., 2011; Damaraju et al., 2014; Lynall et al., 2010).

Dynamic functional network connectivity (dFNC) analyses show that resting-state activity patterns continuously shift between a set of recurring, short-lived connectivity states (Calhoun et al., 2014; Hutchison et al., 2013; Sakoğlu et al., 2010). Alterations in dynamics of the connectivity patterns have been reported in SZ and other disorders (Damaraju et al., 2014; Fu et al., 2025). Patients with SZ show reduced "dynamism", that means fewer state transitions and less variability in connectivity patterns over time (Miller et al., 2016; Rashid et al., 2016). The severity of positive and negative symptoms correlates with fewer transitions between states (Damaraju et al., 2014), suggesting that inflexibility in network dynamics has direct clinical relevance. After antipsychotic treatment, patients more often entered states with stronger inter-network integration, and greater symptom improvement was linked to higher connectivity variability (Cattarinussi et al., 2023).

While the dFNC method offers substantial advantages, analyzing high-dimensional, nonlinear network dynamics poses computational challenges. Information theory provides a powerful framework for quantifying complexity and information processing capacity in neuroimaging data (Poza et al., 2021; Saxe et al., 2018; Yulug et al., 2025). Entropy measures compress high-dimensional data into interpretable scalars that preserve essential dynamics while reducing dimensionality and computational burden, thereby simplifying analysis without sacrificing critical information. Entropy approach has been extensively applied at multiple scales of brain organization from voxel-wise signals (Li et al., 2025; Sokunbi, 2014; Xue et al., 2019) to regional fMRI blood oxygenation level-dependent (BOLD) time series (Nezafati et al., 2020; Saxe et al., 2018; Wang et al., 2014) and to network nodes (Sen et al., 2019; Viol et al., 2017).

In our previous studies (Maksymchuk et al., 2025; Maksymchuk et al., 2024) we introduced dynamic inter-network connectivity entropy (DICE). DICE quantifies the Shannon entropy of how each network distributes its functional connectivity across all other brain networks. Unlike regional entropy approaches that measure signal irregularity, our DICE metric uniquely quantifies the entropy of connectivity patterns. DICE tracks the organizational complexity of connectivity patterns, measuring whether the brain’s inter-network organization becomes more ordered or more random moment-to-moment. High DICE values indicate random distribution of connections across target networks, while low DICE reflects ordered, or organized connectivity patterns. While DICE robustly differentiates SZ from controls, temporal patterns that may capture fundamental aspects of brain pathophysiology remain uninvestigated.

We hypothesize that (1) multiple DICE-derived temporal features capture distinct dimensions of altered network-level dynamics in SZ; and (2) within SZ, distinct DICE-derived dynamic features exhibit different associations with PANSS symptom dimensions, reflecting clinical heterogeneity. To test these hypotheses, we present an analytic framework characterizing DICE time courses using three families of metrics capturing (i) how entropy changes over time (fluctuation magnitude, velocity, acceleration), (ii) the organization of these changes (distributional shape and temporal persistence), and (iii) the repertoire and usage of entropy states (state diversity, occupancy, dwell time). We show that SZ-related alterations in entropy dynamics are multidimensional. Although mean entropy is elevated at the group level, the temporal evolution of entropy varies across networks and individuals and exhibits distinct associations with symptom dimensions, showing that entropy dysregulation is not uniform across patients. Our results indicate that mean entropy alone misses clinically relevant information encoded in the temporal dynamics, which may facilitate biomarker development and treatment targets.

## Methods

### 2.1 Participants, fMRI Data Acquisition and Preprocessing

Resting-state fMRI data was collected from 311 adults, 160 healthy controls (HC) (115 males, 45 females, mean age = 37.03±10.86 years) and 151 individuals with SZ (115 males, 36 females, mean age = 38.77±11.63 years) through the Functional Imaging Biomedical Informatics Research Network (FBIRN) consortium (Potkin & Ford, 2009). SZ diagnoses were confirmed with the structured clinical interview for DSM-IV-TR axis I Disorders (SCID-I/P). Imaging was performed on 3 T systems (Siemens Tim Trio at six sites; GE Discovery MR750 at one site) using gradient-echo EPI (TR/TE = 2 s/30 ms, flip angle = 77°, field of view = 220 mm, 32 ascending axial slices, 4 mm thickness with 1 mm gap). Data preprocessing was performed using the SPM12 toolbox (http://www.fil.ion.ucl.ac.uk/spm/) and included brain extraction, rigid-body head-motion correction, and slice-timing correction to account for timing differences in slice acquisition. The data were then normalized to Montreal Neurological Institute (MNI) template space using an echoplanar imaging (EPI) based registration, resampled to 3 mm³ isotropic voxels, and spatially smoothed with a 6 mm full width at half-maximum Gaussian kernel.

### 2.2 Spatially Constrained Independent Component Analysis

Spatially constrained independent component analysis (ICA) was performed on preprocessed data using GIFT software toolbox http://trendscenter.org/software/gift (Iraji et al., 2021) utilizing the NeuroMark_fMRI_1.0 template (Du et al., 2020). According to this template, 53 ICNs have been categorized into seven functional domains: subcortical (SC), auditory (AUD), sensorimotor (SM), visual (VIS), cognitive control (CC), default mode (DM), and cerebellar (CB). Visualization of the network templates and functional domains is provided in (Du et al., 2020). NeuroMark_fMRI_1.0 provides standardized priors for 53 intrinsic connectivity networks (ICNs), aka functional connectivity networks, derived from large healthy cohorts, enabling spatially constrained ICA that yields harmonized subject-level components (functional networks) across scanners, datasets, and clinical groups. This standardization improves reproducibility, streamlines consistent labeling across functional domains, and supports comparable FNC/dFNC pipelines for cross-study biomarker discovery.

### 2.3 Dynamic Functional Network Connectivity (dFNC)

For each ICN time course, we computed windowed FNC using a sliding-window approach. A tapered sliding window was obtained by convolving a rectangle with a Gaussian (σ = 3 TRs) to partition each ICN time series into several short segments. From the time series for each window, we computed the 53×53 connectivity matrix using pairwise correlations between windowed segments (**Figure 1A**). As a result, for each subject we obtained 53×53×137 tensor that captures FNC changes over time (53 is a number of ICNs; 137 is a number of windows).

**Figure 1.**
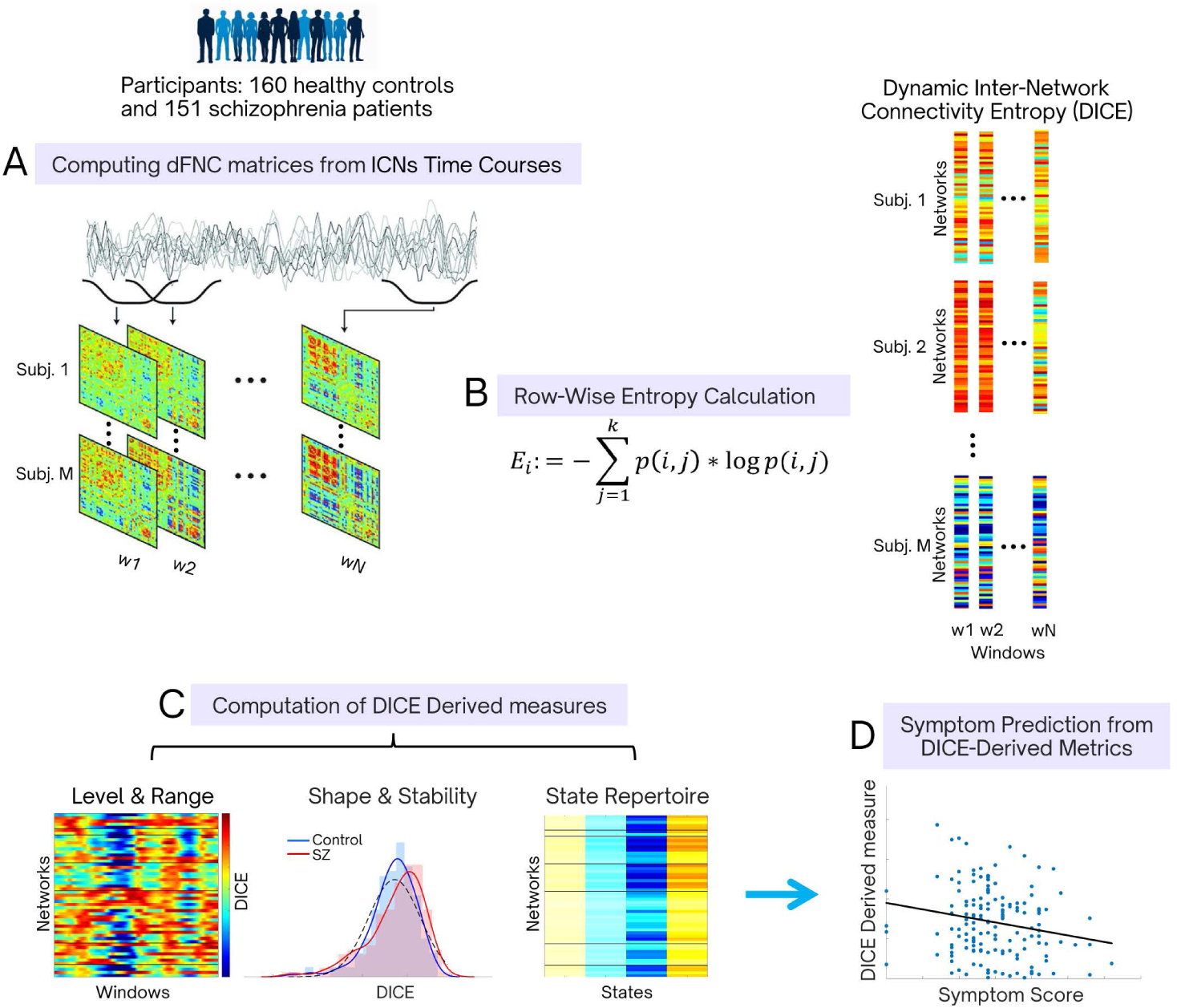
Schematics of main steps of the proposed method. Resting-state fMRI data were decomposed with spatially constrained ICA resulting in 53 ICNs. Based on ICNs time courses, dFNC matrices were computed across all subjects using a sliding-window approach (A). DICE was calculated using dFNC matrices (B). Next, we evaluated three families of DICE derived measures: DICE level and range; DICE shape and temporal organization; state repertoire and occupancy for a centered DICE (C). For each DICE derived measure we examined group differences between SZ and HC as well as associations of each measure with clinical symptoms (D).

### 2.4 Dynamic Inter-network Connectivity Entropy

For each functional network *i*, we used the dFNC matrices to analyze its connectivities with other networks, denoted as *c*(*i*, *j*). These matrices show us how the connectivity strength across different brain networks changes over time. We used *j* ≠ *i*, meaning we examined the connectivity between network *i* and other networks *j*, excluding the connection to itself. To ensure that the connectivity *c*(*i*, *j*) is non-negative, we subtracted the minimum connectivity value, *C*_*min*_ observed globally across all network pairs. This adjustment resulted in a non-negative "translated" connectivity value for each pair, which we denote as *c*^′^(*i*, *j*) = *c*(*i*, *j*) − *C*_*min*_. Next, for each network *i* we computed a probability distribution that consist of a sequence of probabilities {*p*(*i*, 1), *p*(*i*, 2),… *p*(*i*, *k*)}, where *j* ≠ *i*, and each probability *p*(*i*, *j*) is defined as:

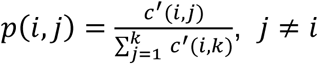

This formula indicates that *p*(*i*, *j*) is calculated by dividing each translated connectivity *c*^′^(*i*, *j*) by the sum of all translated connectivities for network *i* with other networks. Essentially, each *p*(*i*, *j*) represents the proportion of connectivity strength between network *i* and network *j* relative to the total connectivity strength of network *i* to all other networks. Here, *k* represents the total number of networks, which corresponds to the number of possible connections each network could have with others. Next, for every network *i* and window we computed the Shannon entropy, *E*_*i*_, of this probability distribution (**Figure 1B**):

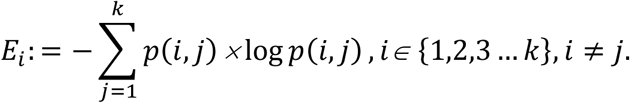

As a result, from 53×53×137 dFNC tensor we obtained 53×137 matrix for each subject. The entropy measures the uncertainty or randomness in the distribution of connectivity strengths for source network *i* across all other networks. A higher entropy value indicates that connectivity strengths are spread more random across networks. In contrast, a lower entropy suggests that network *i* has more concentrated connections with specific networks, indicating more structured connectivity pattern.

This approach provides substantial dimensionality reduction from thousands of pairwise connections to a single entropy vector 53 × 1, where 53 denotes the number of intrinsic functional networks, per time point for each subject. Thus, we obtained tensor in dimensions of 53×137×311, where 137 indicates the number of time windows, and 311 signifies the number of subjects. Such entropy vectors preserve essential information about the temporal evolution of whole-brain connectivity organization.

All computations were performed using custom MATLAB scripts. The implementation of our ICE approach has been incorporated into the GIFT toolbox.

### 2.5 Dynamic Entropy Metrics

We grouped our DICE-based metrics into three families that reflect complementary aspects of ICE dynamics: (1) level and range (baseline entropy, variance, speed and acceleration); (2) shape and temporal organization (distributional shape metrics and zero-crossing rate); (3) state repertoire and occupancy (entropy-state diversity, occupancy and dwell time). The first family captures the entropy baseline, how far it deviates from that baseline, and how quickly and abruptly it changes. The second family evaluates the distributional shape of entropy fluctuations and their temporal structure. The third family captures discrete entropy configurations and how these configurations are used in controls and individuals with SZ.

Our primary analyses focused on group differences and associations between the three entropy-metric families and PANSS subscale scores. More fine-grained network-level item-wise PANSS associations and links with cognitive performance were considered secondary analyses.

#### 2.5.1 Level and Range

Baseline entropy was defined as the temporal mean of the DICE signal across time windows, computed for each network and subject. To capture the overall spread of entropy fluctuations around the baseline, we computed the temporal variance of the DICE signal. To characterize rate of entropy change (velocity) and how abrupt these changes are (acceleration) we computed first and second derivatives of DICE signal, respectively, using numerical gradient estimation with a sampling period of TR = 2 seconds.

#### 2.5.2 Shape and Temporal Organization

For each network and subject, we computed five different distributional measures: absolute skewness (distribution asymmetry measure), excess kurtosis (tail weight measure), composite score (z-scored average of absolute skewness and excess kurtosis, summarizing distributional shape deviations from normality), and Kolmogorov-Smirnov (KS) distance.

Before zero-crossing rate computations, we subtracted temporal mean from DICE timeseries for each network and subject. Zero crossing rate for centered entropy, *Ec*(*t*), was evaluated using a linear interpolation method for sub-sample precision (Gouyon et al., 2000; Kedem, 1986). This approach detects sign changes between consecutive samples and interpolates the exact crossing location assuming linear behavior between samples. Exact zeros were handled by examining the neighborhood context to determine whether they represented true crossings. The zero crossing rate was expressed as crossings per minute, calculated as the total number of crossings divided by the signal duration in minutes.

#### 2.5.3 State Repertoire and Occupancy

We applied fuzzy c-means clustering on centered DICE data to identify distinct entropy configurations. Centered DICE timeseries show relative entropy fluctuations around each subjects’ baseline. It removes individual differences in baseline entropy and focuses on dynamic modulation capabilities. Such approach evaluates true dynamics and individual differences in baseline connectivity don’t affect clustering result. As a result of clustering analysis, we obtained four different entropy states – patterns of entropy deviations from individual baselines.

The number of clusters (k=4) was selected to balance biological interpretability with clustering quality, as determined by silhouette analysis across k=2-10. Clustering was performed with Euclidean distance, using 500 maximum iterations with a minimum improvement threshold of 1×10⁻⁵, replicated across 50 random initializations to ensure stability. The final clustering achieved effective and well-defined separation (partition coefficient = 0.927 is close to 1, and partition entropy = 0.121 is close to 0) while preserving fuzzy membership properties essential for capturing gradual transitions between entropy configurations. The fuzziness parameter of 1.05 provided the best trade-off between cluster compactness and overlap, as indicated by high partition coefficient, low partition entropy, and statistical sensitivity in group comparisons.

For each subject, fuzzy occupancy for each state was computed as the temporal average membership probability from the fuzzy partition matrix. Dwell time was derived from a hard state sequence obtained by assigning each window to the state with the maximum membership. For each state, we identified all consecutive runs and defined dwell time as the mean run length in windows. If a state was never visited, its dwell time was set to NaN.

Also, for each subject, we computed state membership entropy at every window as the Shannon entropy of the fuzzy state memberships and then averaged these values across windows. The membership entropy of entropy states, we will call entropy-state diversity for simplicity.

### 2.6 Connecting Group-Level Statistics to Spatial ICA Maps

All image processing and analysis were implemented in MATLAB R2024a (MathWorks). We used spatial ICA maps of 53 ICNs from the Neuromark1 fMRI 1.0 template (Du et al., 2020) with 4D volume of dimensions x×y×z×k that was reshaped into a two-dimensional matrix of voxels×networks by collapsing the spatial dimensions. To restrict analysis to brain voxels, we computed the row-wise sum of absolute values across all components and retained only those voxels with nonzero loading. Each component was divided by its maximum absolute value across all voxels. To incorporate group differences, each normalized network was multiplied by its corresponding T-value, producing a weighted matrix. Then we collapsed a weighted matrix back to a single composite map by taking the voxel-wise maximum across components, so that each voxel’s value directly reflects the largest t-statistic among all networks. To generate multi-slice montages of our volume overlaid on a standard anatomical template we used the icatb_image_viewer function from the GIFT toolbox (Iraji et al., 2021) (http://trendscenter.org/software/gift. All the panels in those montages are axial slices, arranged in a grid so that each tile shows the brain at a different z-coordinate (from −53 mm up to +80 mm in 9 mm steps). Applying a threshold=2 helps focus on voxels where the effect is not just statistically significant at the network level, but also strongly expressed spatially.

### 2.7 Diagnosis Effect

To estimate the effect of diagnosis on the measures investigated here we applied multiple linear regression analysis including age, sex, gender, and head motion (MFD) as covariates. For each network *k*, we used the following model:

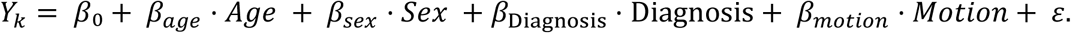

Were *Y*_*k*_ is a DICE-derived measure, the diagnosis is binary (SZ = 1, HC = 0), motion is a mean frame displacement (MFD). Accordingly, *β*_Diagnosis_ reflects the mean difference (SZ–HC), *β*_Diagnosis_ > 0 indicates higher values in SZ, while *β*_Diagnosis_ < 0 indicates lower values in SZ. We controlled the false discovery rate (FDR) across the 53 networks within each measure using the Benjamini–Hochberg (BH) procedure at α=0.05.

### 2.8 Associations of DICE-derived Measures with Symptom Severity

Clinical symptom severity was assessed using the Positive and Negative Syndrome Scale (PANSS). Associations between DICE-derived measures and symptom severity were evaluated at both the PANSS subscale and item-wise levels, at whole brain and network levels. Only SZ patients were included in this analysis. For PANSS subscale analyses at network level we fitted separate regression model for each brain network *k*:

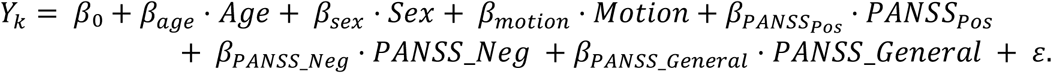

For item-level analyses network level, we fitted separate models for each of the 30 PANSS items and each network *k*:

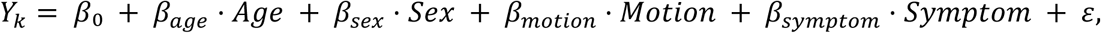

where symptom is one of thirty PANSS item.

For network-level analyses at both the PANSS subscale and item levels, p-values were corrected for multiple comparisons across the 53 networks using the Benjamini–Hochberg FDR procedure, α=0.05.

For whole-brain analyses, each DICE-derived measure was averaged across the 53 networks within each subject, and the resulting global values were analyzed using regression models analogous to the network-level analyses.

## Results

### 3.1 Entropy Level and Range

#### 3.1.1 Elevated Baseline Entropy and Reduced Dynamic Range

Using covariate adjusted regression model, we found that out of 53 brain networks, 41 (77%) showed significantly higher temporal mean DICE (**Figure 2A**), and 48 of 53 (90.6%) networks showed significantly lower DICE temporal variance in SZ (**Figure S1A**). The covariate-adjusted group difference for DICE variance was also highly significant at the global level (p = 5.94×10^-5^).

**Figure 2.**
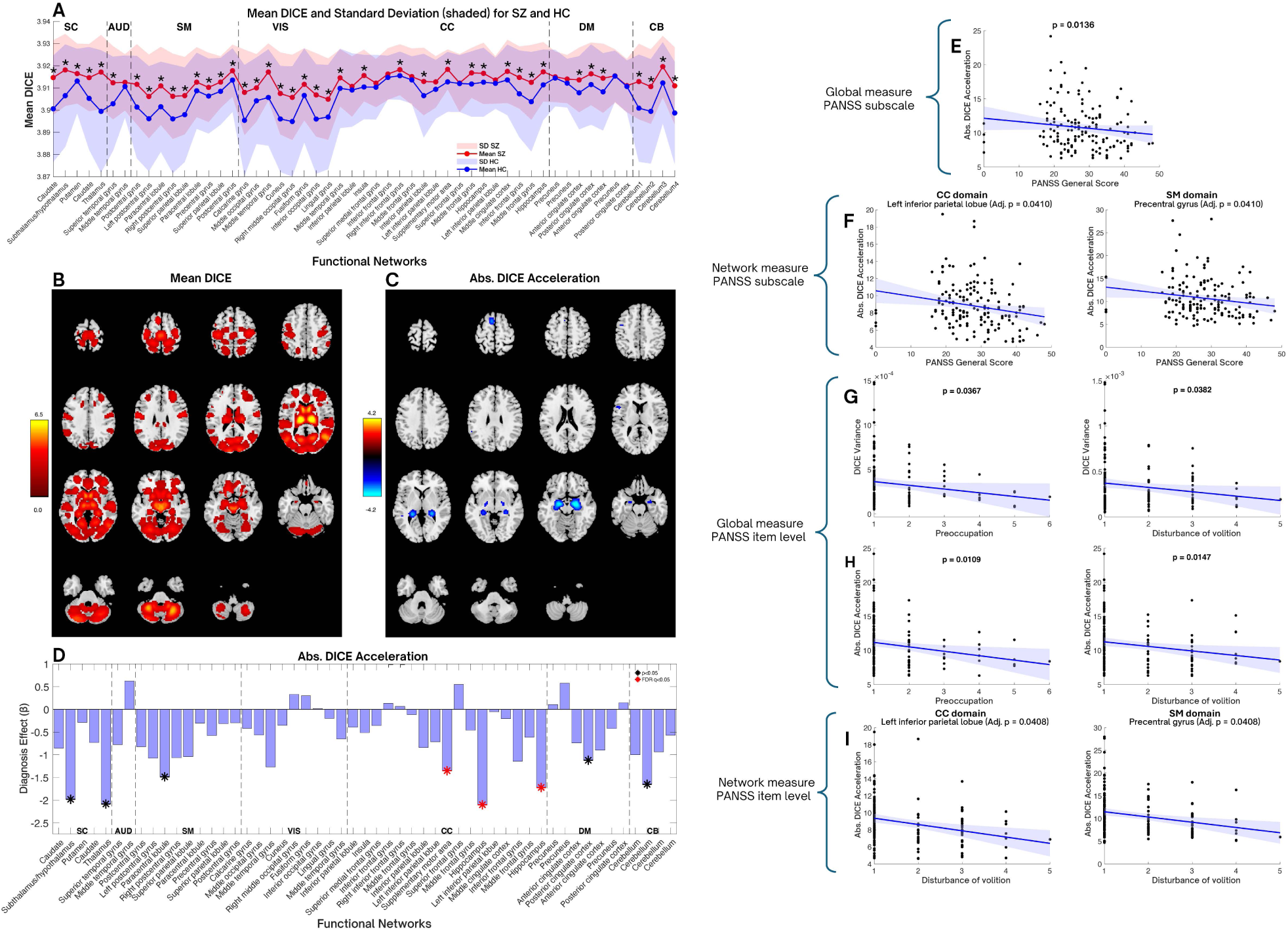
Diagnosis effects and PANSS associations and for level and range metrics. (A) Mean DICE and SD for SZ (blue) and HC (red). Asterisks indicate networks with statistically significant group differences in entropy, all survived FDR correction at α=0.05 (B) SZ-HC group differences (T-values) in mean DICE (B) and acceleration (C), displayed on spatial ICA maps. The red areas indicate SZ>HC, whereas blue areas show SZ<HC. Axial brain slices arranged in a grid showing the brain at a different Z-coordinate (from –53 mm up to +80 mm in 9 mm steps). (D) Diagnosis effect, *β*, for entropy acceleration. Black asterisks show p-values less than 0.05, red asterisks indicate p-values survived FDR correction; positive HC>SZ. (E-F) Reduced acceleration associates with PANSS general subscale at whole-brain and network levels (CC, SM domains). (G-H) Both variance and acceleration metrics converge on specific symptoms at the global level, particularly disturbance of volition and preoccupation. (I) Network-level analyses confirm symptom associations in cognitive control and sensorimotor regions. Solid lines indicate fitted regression trends with shaded 95% confidence intervals. All results are obtained using regression analysis controlling for confounds (age, sex, and head motion). Adjusted p-values reflect BH FDR correction across the 53 networks.

We connected group-level statistics (t-values of entropy differences) to spatial ICA maps and displayed these group-level differences on brain slices (**Figures 2B, S1B).** These are T-values from group comparisons, with FDR correction with BH procedure. Entropy increase and variance decrease are remarkably widespread across brain networks in SZ, indicating the pervasive nature of connectivity disorganization. Extensive changes were observed across multiple neuroanatomical levels. Lower slices clearly show impairment in the cerebellum, crucial for cognitive coordination. Middle slices show pronounced effects in the subcortical and auditory regions. Upper slices show widespread cortical impairments including frontal, parietal and sensory regions, that is consistent with executive, attention, integration and sensory processing deficits in SZ.

Altered entropy and entropy temporal variance in SZ has a global nature, it is not a focal damage, it is brain-wide changes in functional connectivity organization and temporal dynamics of this organization. Mean entropy is consistently higher and variance is consistently lower for every region that show statistically significant group difference. The most intense effects occur at critical information hubs: thalamus, subthalamus, caudate and cerebellum. Thus, SZ involves a global shift in how brain networks organize their connectivity, not isolated to one system but affecting the entire brain’s information routing architecture.

#### 3.1.2 Suppressed Second-Order Entropy Dynamics in Cognitive Control Networks

Next, we evaluated covariate-adjusted group differences in first- and second-order entropy dynamics, velocity, *dE*/*dt*, and *d*^2^*E*/*dt*^2^, by estimating the diagnosis effect in regression models with age, sex, and head motion as covariates. No effects of diagnosis on entropy velocity survived FDR correction. In contrast, three networks in the CC domain survived FDR correction for absolute entropy acceleration: the supplementary motor area and two hippocampal networks (**Figure 2D**). The corresponding spatial ICA maps for the brain networks that survived FDR correction are shown in **Figure 2C**. The negative diagnosis coefficient, *β*, indicates higher absolute acceleration in healthy controls relative to patients. Additional networks in SC, SM, DM, and CB domains showed uncorrected diagnosis associations, but none survived FDR correction across 53 networks (**Figure 2D**).

#### 3.1.3 Associations with PANSS scores

Level and range metrics showed convergent associations with clinical symptoms. Both reduced DICE variance and DICE absolute acceleration are associated with greater PANSS general severity at the global level, particularly for disturbance of volition and preoccupation (**Figure 2G,H**). The DICE acceleration–symptom relationship shows remarkable consistency across levels and scales. The negative association emerges at the whole-brain level (**Figure 2E**) and across CC and SM networks (left inferior parietal lobule and precentral gyrus, **Figure 2F**), extends to additional CC regions (inferior parietal lobule, middle frontal gyrus, hippocampus) and the DM (anterior cingulate cortex) (**Figure S2**), and is observed in both PANSS subscale (**Figure 2E,F**) and item-level analyses (**Figure 2H,I**). This convergence across independent metrics, spatial scales, and symptom measures indicates that impaired entropy modulation, both in DICE magnitude (variance) and DICE acceleration may represent a core SZ deficit in dynamic flexibility associated with volitional and cognitive symptom severity.

### 3.2 Shape and Temporal Organization

#### 3.2.1 SZ Cognitive Control and Cerebellar DICE Distributions are More Gaussian than Controls

Next, we computed distinct DICE distributional characteristics for both groups and evaluated covariate adjusted group comparisons for every measure. Two networks survived multiple comparisons correction for KS distance: middle frontal gyrus (network 31, CC domain) and cerebellum (network 52, CB domain, **Figures 3A and S3A**) and one network for composite score: inferior frontal gyrus (network 29, CC domain, **Figures 3B and S3B**). Although absolute skewness and the excess kurtosis showed significant differences in the CC and CB domains, no networks remained significant after multiple comparisons correction (**Figure S3C-D**).

**Figure 3.**
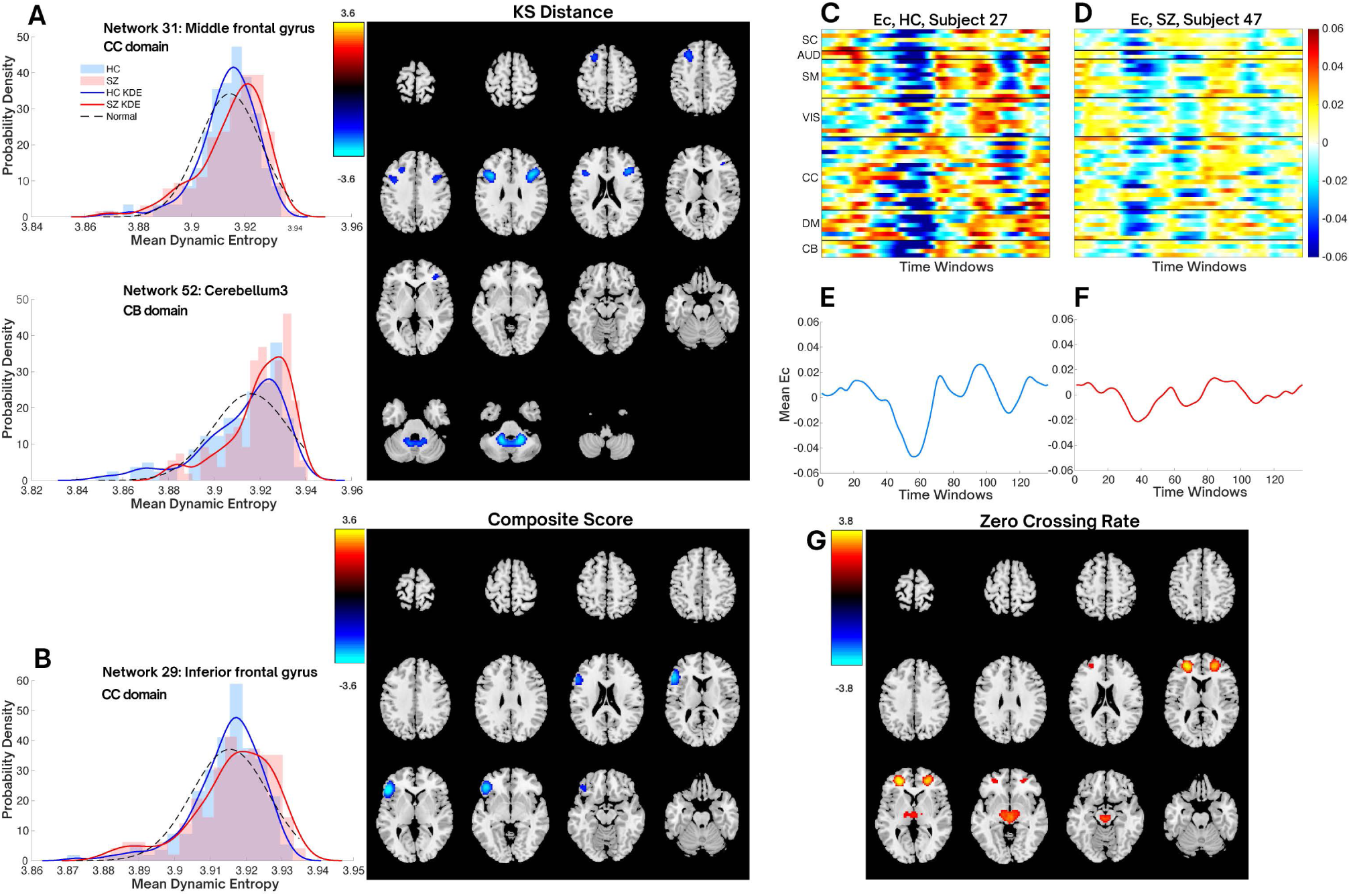
Diagnosis effects for entropy distributional measures and zero crossing rate. (A) Representative subject-level group distributions with kernel density (KD) estimates for networks (middle frontal gyrus and cerebellum3) showing a significant diagnosis effect on KS distance after multiple-comparison correction. Right panels show spatial ICA maps for these networks, illustrating reduced KS distance in SZ. (B) Representative subject-level group distributions with kernel density (KD) estimates for inferior frontal gyrus showing a significant diagnosis effect on the composite score after multiple-comparison correction. Right panels show spatial ICA maps for inferior frontal gyrus. (C-D) Colormaps showing temporal fluctuations of centered DICE across 53 functional networks for two presentative subjects, control and SZ. (E-F) Fluctuation of mean centered DICE averaged across 53 functional networks for two representative subjects: control and SZ. (G) Covariate adjusted group differences in mean zero-crossing rate (T-values), displayed on spatial ICA maps; the color bar denotes T-values. Axial brain slices arranged in a grid showing the brain at a different Z-coordinate. All results are obtained using regression analysis controlling for confounds (age, sex, and head motion).

These results show that relative to HC, individuals with SZ exhibit not only higher mean entropy, but also lower distance from normality of their entropy distributions for CC and CB networks. The consistency of the direction (HC group has larger distance from normality) across these networks and across multiple distributional measures (**Figure S3**) provide strong evidence about reduced healthy dynamic complexity in SZ.

#### 3.2.2 Reduced Zero Crossing Rate in Cognitive Control and Subcortical Networks in SZ

Temporal evolution of centered DICE, *Ec*(*t*), across all functional domains is presented in **Figure 3 C-D** for two representative subjects, SZ and control. *Ec*(*t*) fluctuates around zero, taking positive values when entropy is above baseline and negative values when it is below baseline (**Figure 3E-F**). **Figure S6A** illustrates that zero-crossing rate is significantly higher across networks in SC, SM, VIS, CC and CB domains. However, only two networks survived multiple comparisons correction: subthalamus/hypothalamus (SC domain) and middle frontal gyrus (CC domain). These networks are shown on spatial ICA maps (**Figure 3G**). Higher zero crossing rates in SZ patients means reduced temporal persistence of above/below mean entropy excursions.

#### 3.2.3 Associations with PANSS scores

A shift toward more Gaussian entropy dynamics in SZ is linked to increased PANSS positive symptom severity. Specifically, reduced composite scores predicts higher PANSS positive subscale scores both globally and in middle cingulate cortex (MCC), **Figure 4A,B** respectively. PANSS item-level analyses confirmed associations with delusions and hallucinatory behavior (**Figure 4C**). Similarly, reduced KS distance associates with positive symptoms including delusions, hallucinatory behavior, and conceptual disorganization (**Figure 4D**). In addition, item-level analyses showed associations with general symptoms like unusual thoughts (both for composite score and KS distance), disturbance of volition, anxiety and disorientation (**Figure S4**). Individual distributional components (skewness, kurtosis) showed convergent patterns. Reduced absolute skewness associated with increased PANSS positive scores (**Figure S5A**) in MCC. PANSS item-level analyses confirmed absolute skewness associations with delusions and hallucinatory behavior (**Figure S5B**), while reduced kurtosis predicted delusions and unusual thoughts (**Figure S5C**).

**Figure 4.**
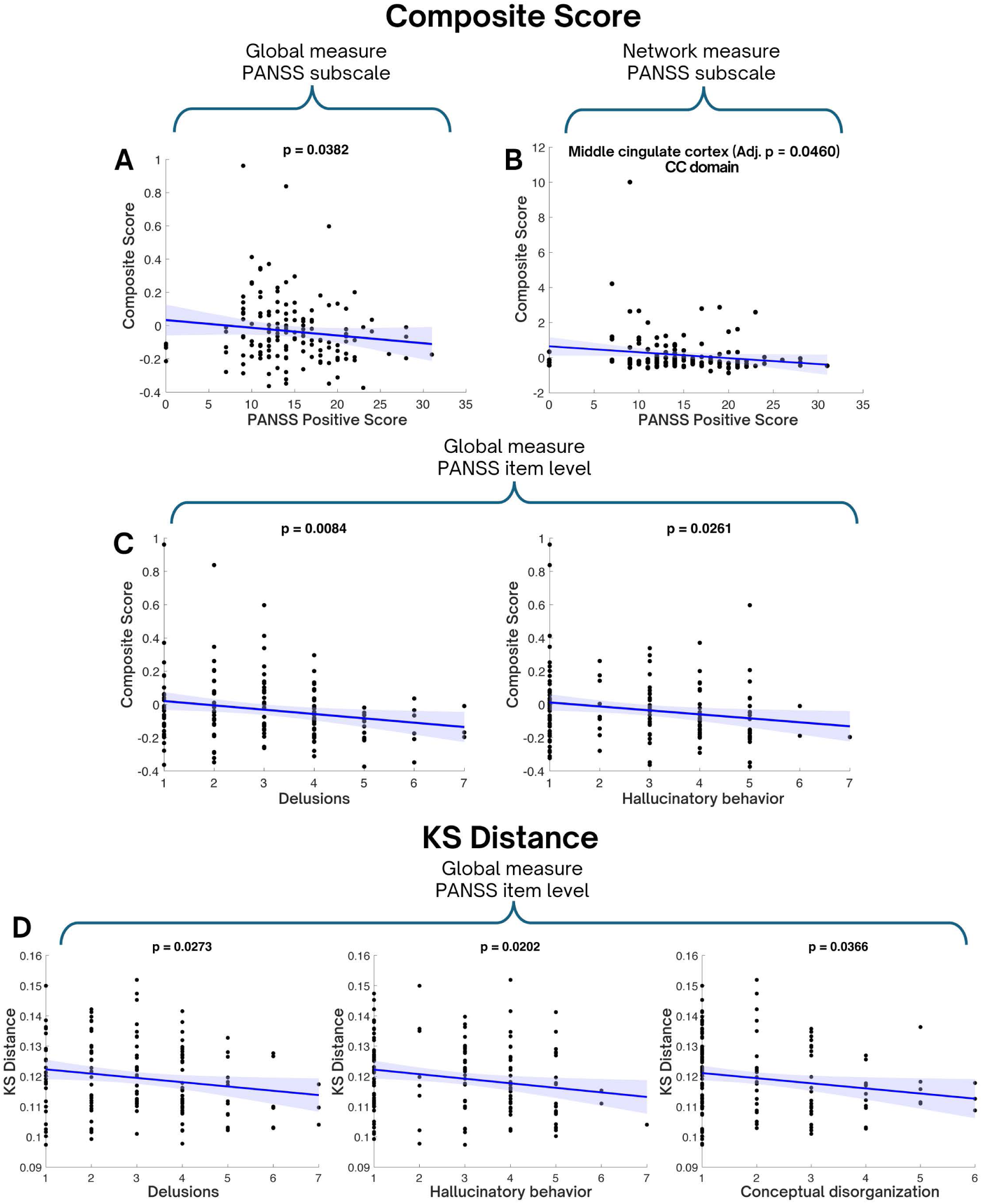
Convergent distributional metrics (composite score and KS distance) are associated with positive symptom severity across scales in SZ: PANSS subscale - Composite score associations at global level (A) and network level (B); Global item-wise PANSS associations for composite score (C) and for KS distance (D). Solid lines indicate fitted regression trends with shaded 95% confidence intervals. All associations were evaluated with linear regression approach controlling for age, sex, and head motion. Adjusted p-values reflect BH FDR correction across the 53 networks.

Higher zero crossing rate (lower temporal persistence) for *Ec*(*t*) within patients is associated with greater symptom severity across all PANSS domains. Globally, higher zero crossing rate associated with more severe hallucinatory behavior, disturbance of volition, and disorientation (**Figure S6B**). Network-level analyses revealed further symptom associations (**Figure S6C**), spanning positive symptoms, including hallucinatory behavior and conceptual disorganization (mapped to the left postcentral gyrus and left inferior parietal lobule) and negative symptoms, including stereotyped thinking and emotional withdrawal (mapped to the left inferior parietal lobule and precuneus). These finding demonstrate that temporal instability impacts multiple symptom dimensions.

### 3.3 State Repertoire and Occupancy

#### 3.3.1 Reduced Entropy-State Repertoire and Shifted State Occupancy in SZ

Fuzzy clustering on centered DICE data identified four cluster centroids with distinct entropy states, patterns of entropy deviations from individual baselines across networks (**Figure 5A**). Patients showed significantly reduced entropy-state diversity of (p=0.004) (**Figure 5B**). Lower entropy-state diversity indicates that patients are assigned to specific states more deterministically, reflecting reduced flexibility in modulating entropy around their individual means. In contrast, the control group shows fuzzier, more flexible state transitions characteristic of healthy brain dynamics.

**Figure 5.**
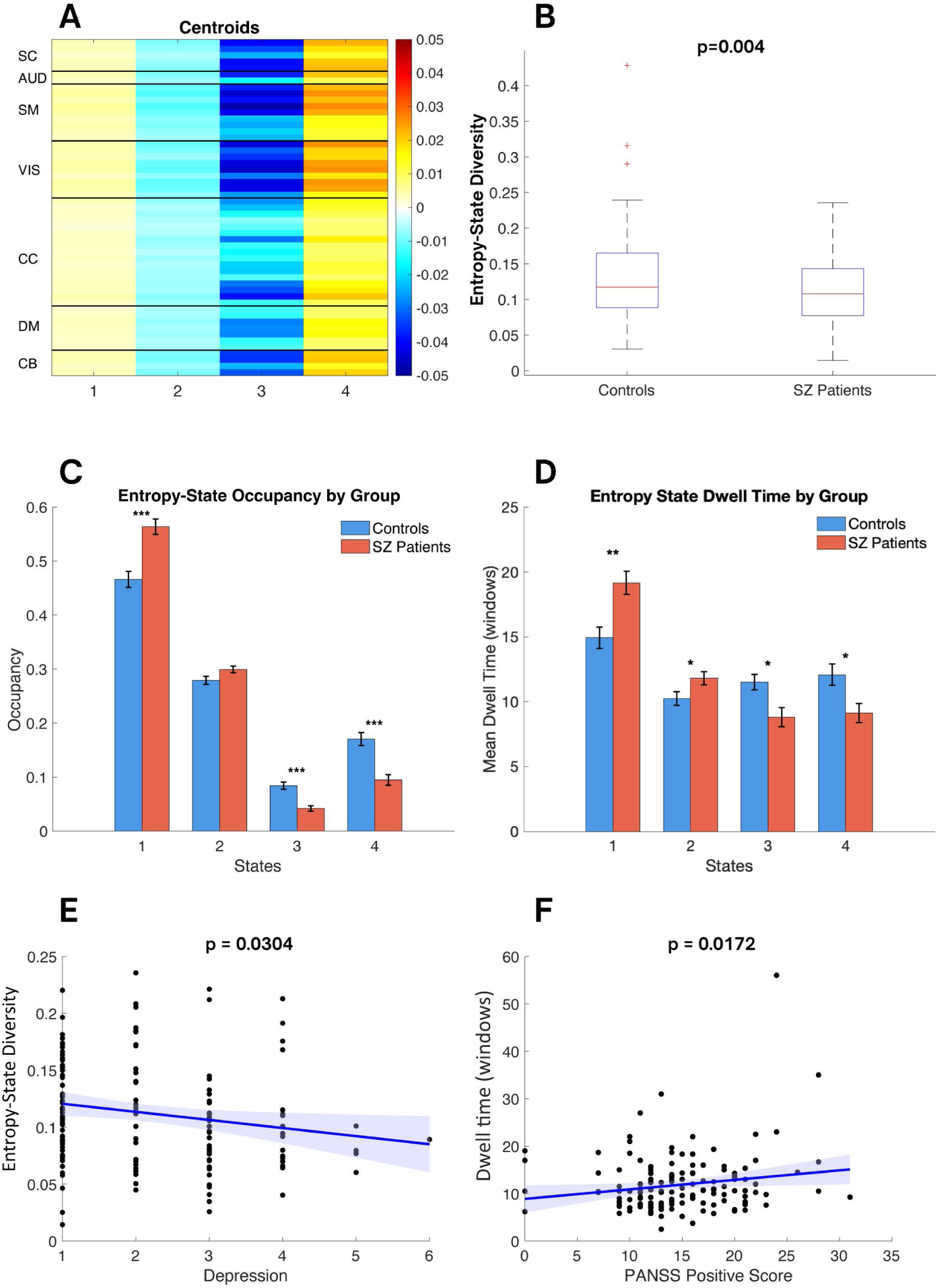
Patients with SZ show altered centered DICE dynamics and reduced entropy-state diversity relative to healthy controls. (A) Fuzzy c-means clustering produces four recurrent patterns of centered DICE: two near-baseline entropy patterns (states 1, 2), and patterns with high entropy deviations (states 3, 4). (B) Entropy-state diversity for SZ and controls. Mean fuzzy occupancy (C) and dwell time (D) per state. (E) Association of reduced state diversity with higher depression score. (F) Prolonged dwell time in near-baseline state 2 linked to higher PANSS symptom scores. All group differences and symptoms associations were evaluated using linear regression controlling for age, sex, and head motion. Solid lines indicate fitted regression trends with shaded 95% confidence intervals.

Patients with SZ demonstrated altered occupancy in three of four entropy states and dwell time differences in all configurations, FDR p<0.05 (**Figure 5C-D**). Patients spend more time and stay longer in states where entropy deviations are small (states 1,2), while control group preferentially occupies and dwells in extreme states 3 and 4 with high deviations from mean entropy (states 3, 4). These effects were highly significant after FDR correction (**Table 1**). This is consistent with our previous findings related to reduced magnitudes of entropy fluctuations around mean for SZ patients.

**Table 1.** Diagnosis effect (β) and p-values for fuzzy occupancy and dwell time across four states of centered DICE controlling for age, sex, and head motion. BH FDR significant p-values are shown in bold.

| DICE Clusters | | $\beta$ diagnosis | p-values |
| --- | --- | --- | --- |
| Fuzzy Occupancy | State 1 | 0.0960 | **$1.084 \times 10^{-5}$** |
| | State 2 | 0.0179 | $7.963 \times 10^{-2}$ |
|  | State 3 | -0.0391 | **$1.335 \times 10^{-5}$** |
|  | State 4 | -0.0748 | **$6.116 \times 10^{-6}$** |
| Dwell Time | State 1 | 4.5275 | **$4.450 \times 10^{-4}$** |
|  | State 2 | 1.5683 | **$3.704 \times 10^{-2}$** |
|  | State 3 | -2.7054 | **$6.244 \times 10^{-3}$** |
|  | State 4 | -2.7181 | **$2.644 \times 10^{-2}$** |

Notably, that entropy can be near baseline across all networks (states 1, 2) and highly deviate from mean across majority of networks in SC, SM, VIS and CB domains (states 4,5), reflecting wider entropy ranges for these networks. This finding is consistent with higher variance of non-centered entropy in these domains.

#### 3.3.2 Associations with PANSS scores

Reduced state diversity associated with greater depression severity (**Figure 5E**). Prolonged dwell time in near-baseline entropy state 2 is associates with higher PANSS positive scores (**Figure 5F**). Item-wise analysis confirmed positive associations of longer dwell time in near baseline entropy states (1 and 2) with positive symptoms such as grandiosity, delusions, and unusual thoughts (**Figure S7, A-D**), indicating that inability to exit near-baseline entropy states underlies positive symptom expression. Increased occupancy in near baseline entropy state 1 and reduced occupancy in high-deviation states (3 and 4) are associated with preoccupation (**Figure S7, E-G**).

## Discussion

Temporal fluctuations of DICE capture the intrinsic "breathing" of brain network organization, alternating between organized (low entropy) and disorganized (high entropy) inter-network connectivity. Our analysis reveals that this neural breathing in SZ exhibits "rigid randomness" characterized by: (1) globally elevated entropy with reduced fluctuation magnitude and diminished acceleration; (2) shifted toward Gaussian entropy distributions and reduced temporal persistence in CC, SC and CB networks; (3) restricted entropy-state repertoire with patients trapped in near-baseline states and rarely visiting states with large entropy deviations; (4) this multidimensional disruption in SZ map to distinct symptom profiles, reflecting clinical heterogeneity of SZ. All these findings extend beyond traditional dysconnectivity studies (Friston & Frith, 1995; Stephan et al., 2006; Stephan et al., 2009; Weinberger, 1992), indicating that SZ involves global reconfiguration of how brain networks organize their connectivity patterns over time.

### 4.1. Elevated Randomness with Reduced Dynamic Range

Elevated baseline entropy together with reduced variance and entropy acceleration in patients indicates that inter-network connectivity organization is more random, while the temporal reconfiguration of this organization is less variable and less abruptly changing over time. This pattern is consistent with computational models emphasizing reduced signal-to-noise ratio arising from disrupted synaptic and neuromodulatory control in SZ (Rolls et al., 2008). More broadly, NMDA receptor hypofunction has been proposed as a key mechanism in SZ implicating aberrant large-scale network interactions and impaired dynamic regulation of cortical network activity (Nakazawa & Sapkota, 2020).

The global elevation of baseline entropy, together with low-magnitude fluctuations, provides a quantitative characterization of the reduced connectivity dynamics consistently reported in SZ and supports the view that dysconnectivity reflects widespread network disruption (Damaraju et al., 2014; Miller et al., 2016; Rashid et al., 2016; Stephan et al., 2009). In this framework, small fluctuations around an elevated baseline indicate a restricted dynamic range of inter-network connectivity and an impaired ability to dynamically adjust interactions between networks.

Moreover, simultaneously increased baseline randomness in connectivity strength distribution with decreased dynamic complexity explains seemingly contradictory hypo-and hyper-connectivity findings in SZ literature. Higher entropy yields a diffuse, poorly selective coupling landscape that inflates weak correlations between networks (apparent hyperconnectivity), while suppressed dynamics limit excursions into strongly organized states, weakening specific connectivity patterns (hypoconnectivity). Which pattern emerges depends on network examined and analytic scale.

Although SZ shows reduced dynamism at large-scale functional network level, analyses at fine spatial (voxel, single networks) and temporal (TR-wise) scales reveal increased signal "wildness". Patients with SZ have higher power at high temporal frequencies across multiple spatial scales (Miller et al., 2015), increased prevalence of transient spectral peaks in network time courses at TR-level resolution (Miller & Calhoun, 2020), and elevated spatiotemporal derivatives at the voxel level (Miller et al., 2023). Yet coarse-grained sliding window analyses at the functional network level reverse the picture: patients become stuck in restricted regions of the state space (Miller et al., 2016). Our entropy framework captures both aspects of this scale-dependent phenomenon.

The elevated baseline entropy reflects accumulated effects of fine-scale disorder, while reduced dynamic range and decreased acceleration reveal the emergent rigidity at the network level. This isn’t paradoxical but rather reflects how spatial aggregation and temporal windowing transform microscale noise into macroscale inflexibility.

### 4.2 More Gaussian Temporal Dynamics

Our findings extend existing works on dynamic functional connectivity (Calhoun et al., 2014; Fu et al., 2019; Miller et al., 2016; Sakoğlu et al., 2010) by providing an information-theoretic framework quantifying how the organizational complexity of connectivity patterns evolves over time. Whereas previous studies identified discrete connectivity states showing reduced state transitions, longer dwell times in weakly connected configurations, and reduced state-space trajectory complexity in SZ, our DICE framework reveals the distributional properties underlying this rigidity. We demonstrate that the reduced flexibility observed in discrete clustering reflects a shift toward less skewed, less extreme and more Gaussian temporal dynamics, where elevated baseline entropy paradoxically coexists with a restricted repertoire of connectivity configurations.

Our empirical findings of a shift toward more Gaussian entropy distributions align with theoretical work demonstrating that heavy-tailed connectivity distributions support richer neural dynamics (Xie et al., 2025). In their models, heavier-tailed connectivity supports a broad edge-of-chaos regime at the cost of compressed, lower-dimensional attractors, a tradeoff that echoes our finding that more Gaussian-like entropy distributions in SZ accompany elevated baseline entropy and reduced dynamical state diversity. This pattern also consistent with the broader dynamical systems perspective reported by (Breakspear, 2017), who demonstrated that healthy cortical activity exhibits non-Gaussian, metastable dynamics, while psychiatric conditions involve disturbances restricting state-space exploration. Together, these theoretical frameworks support our SZ findings reflecting a shift away from a metastable, dynamically rich regime toward a more Gaussian, noise-like regime with higher baseline entropy but suppressed dynamic complexity.

### 4.3. Reduced State Repertoire and Prolonged Dwell-times in Near Baseline Entropy States

Reduced entropy-state diversity and prolonged dwell-times in states with decreased entropy fluctuations align with reports of suppressed connectivity variability and diminished connectivity dynamism in SZ patients (Blair et al., 2024; Damaraju et al., 2014; Miller et al., 2016; Puvogel et al., 2022). However, our DICE-based framework clarifies the underlying mechanism, the reduction is not simply in the number of state transitions but in the capacity to modulate connectivity organization away from baseline configurations.

### 4.4 Clinical Heterogeneity Reflected in Entropy Dynamics

Our DICE-based framework revealed that alterations in entropy dynamics are tightly coupled to symptom severity in SZ. Different entropy metrics predict distinct symptom dimensions. Reduced DICE variance and acceleration are associated with higher general symptom severity, particularly disturbance of volition and preoccupation, indicating that patients with the most “rigid” entropy dynamics exhibit more severe motivational deficits and more inflexible mental focus. This supports the idea that impaired dynamical flexibility in functional connectivity patterns contributes to clinical inflexibility (Cattarinussi et al., 2023; Damaraju et al., 2014; Miller et al., 2016).

Reduced deviation from normality at global and network level, especially in middle cingulate cortex (MCC), is associated with higher PANSS positive scores, particularly delusions and hallucinations. This aligns with established roles of the MCC in salience processing and error monitoring. As a salience network hub, MCC dysfunction disrupts the detection and evaluation of behaviorally relevant stimuli, leading to aberrant salience attribution that underlies delusions and hallucinations (Jardri et al., 2011; Palaniyappan & Liddle, 2012; Uddin, 2015).

Temporal persistence of entropy oscillations shows broad symptom associations. Globally, reduced temporal persistence (higher zero-crossing rate) is associated with more severe hallucinatory behavior, volition disturbance and disorientation. At the network lever reduced temporal persistence in inferior parietal lobule, precuneus, and left postcentral gyrus is associated with greater conceptual disorganization, stereotyped thinking, and emotional withdrawal. The inferior parietal lobule and precuneus overlap the canonical DM and fronto-parietal control systems, whose abnormal connectivity has been implicated in thought disorder, positive, and negative symptoms (Sendi et al., 2021; Wang et al., 2015). A link between the left postcentral gyrus and auditory hallucinations in SZ was previously reported in (Nenadic et al., 2010; Zaykova et al., 2025) primarily involving gray matter volume reductions and aberrant functional connectivity.

Reduced entropy-state diversity is associated with higher depression scores, consistent with metastability models of SZ linking a restricted state repertoire and fewer transitions to reduced behavioral flexibility and affective flattening (Damaraju et al., 2014; Miller et al., 2016). Longer dwell time in near-baseline entropy states relates to greater positive symptom severity (grandiosity and delusions), whereas reduced occupancy of high-deviation states relates to greater general psychopathology (preoccupation). Prior dFNC studies consistently report that prolonged residence in weakly integrated connectivity states and reduced transitions into strongly integrated configurations are linked to more severe symptoms and poorer treatment outcomes (Cattarinussi et al., 2023; Damaraju et al., 2014; Zhang et al., 2021).

### 4.5 Limitations and Future Directions

Several limitations should be considered. First, while our cross-sectional design reveals robust group differences, longitudinal studies are needed to determine whether entropy alterations represent trait markers. Following patients through different illness phases and treatment responses could clarify the stability and clinical utility of entropy measures.

Second, medication effects remain a potential confound. Although entropy alterations reflect intrinsic pathophysiology rather than medication effects alone (given their widespread nature and specific symptom associations), future studies should examine medication-naive first-episode patients and systematically evaluate how different antipsychotics modulate entropy dynamics.

Third, what we observe at the network level might differ substantially from dynamics at the voxel level. Our network-level entropy measures capture emergent properties that may obscure important fine-scale phenomena. Future studies should explicitly examine entropy dynamics across multiple spatial and temporal scales to fully characterize the scale-dependent nature of neural alterations.

### 4.6 Conclusion

Our DICE framework enables substantial whole-brain dimensionality reduction while preserving essential dynamic information, yielding both computational efficiency and interpretability. We revealed that SZ has multidimensional alterations of entropy dynamics. Within SZ patient different aspects of entropy dynamics relate to distinct symptom profiles, revealing a rich symptom–network mappings and highlighting the heterogeneity of SZ. These findings advance our understanding of SZ as a disorder of multiple, dissociable abnormalities in entropy dynamics, providing anatomically specific and clinically relevant targets for developing precision biomarkers and guiding symptom-focused interventions.

