## Supplementary Materials for "When Randomness Becomes Rigid: Dynamic Connectivity Entropy and Symptom-Linked Network Dysfunction in Schizophrenia"

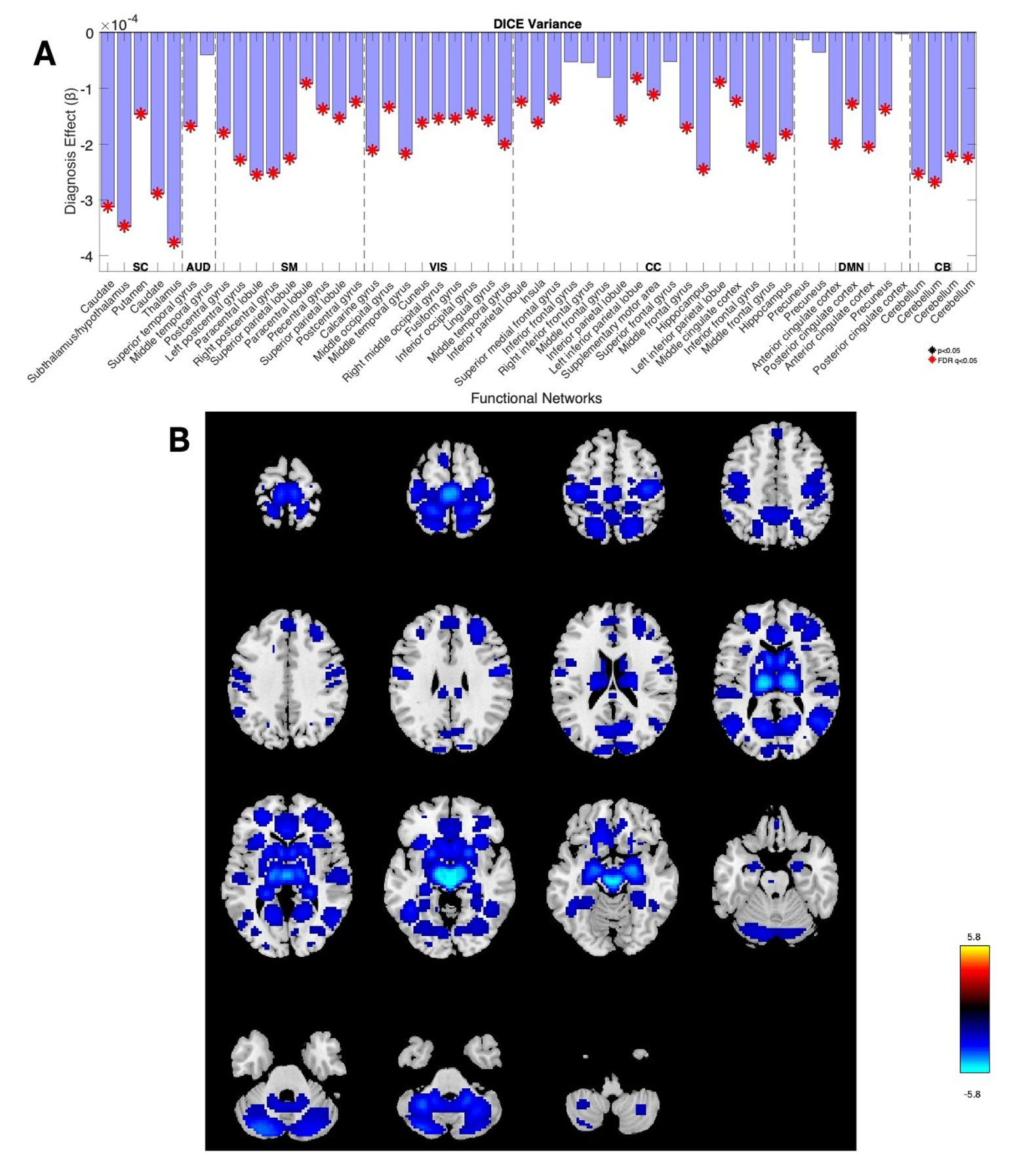

**Figure S1.** SZ patients have reduced DICE temporal variance relatively to controls across majority of brain networks. (A) Diagnosis effect, $\beta,$for DICE variance. Red asterisks indicate p-values survived FDR correction across 53 networks at $\alpha=0.05$; β<0 indicates lower values in SZ. (B) Covariate adjusted SZ-HC group differences (T-values) in DICE variance displayed on spatial ICA maps. Axial brain slices arranged in a grid show the brain at a different Z-coordinate (from –53 mm up to +80 mm in 9 mm steps).

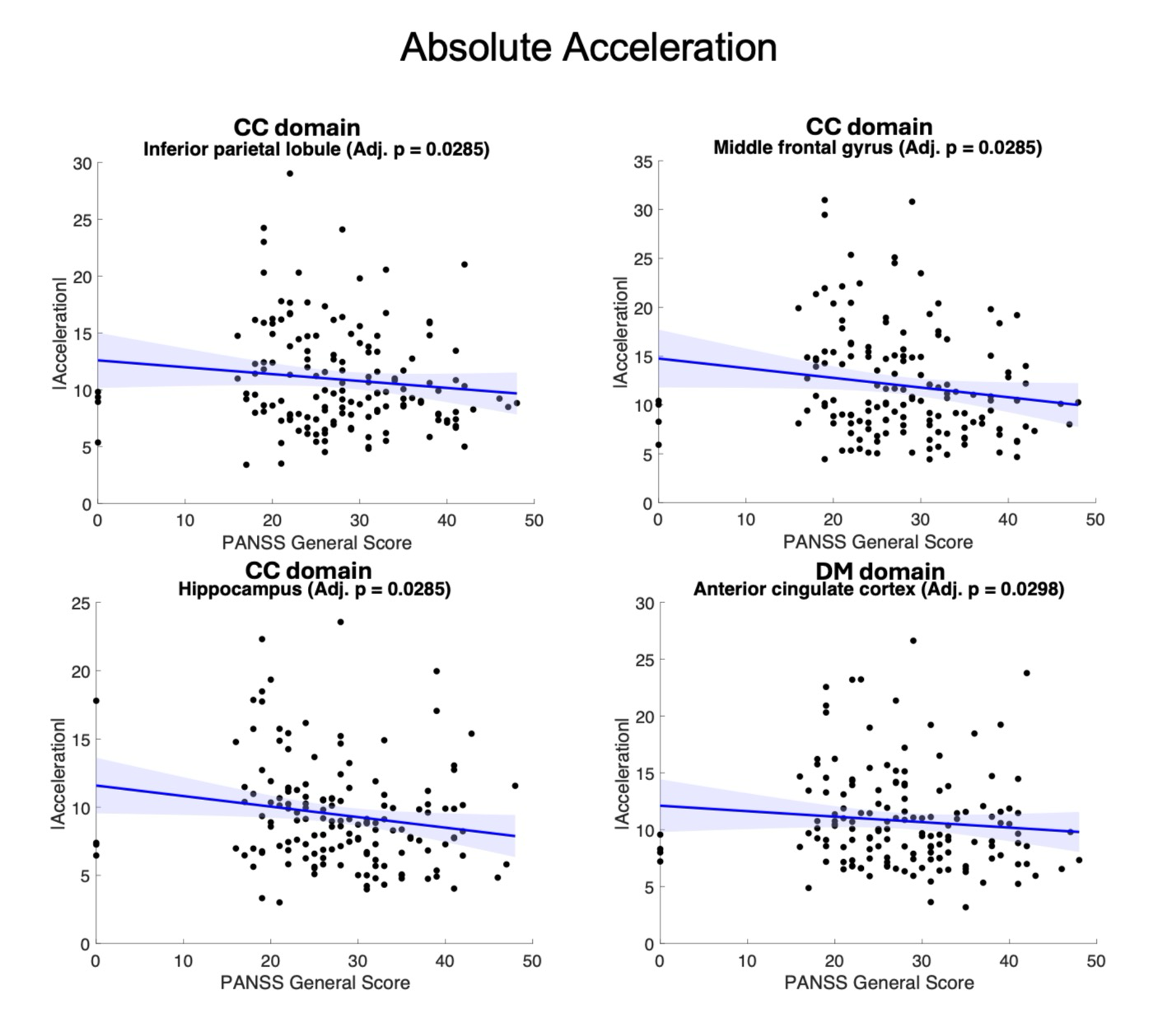

**Figure S2 (Supplementary to Figure 2).** Associations between network-level DICE absolute acceleration and item-vise PANSS symptoms. Solid lines show fitted linear regression trends; shaded bands indicate 95% confidence intervals. Associations were tested using linear regression controlling for age, sex, and head motion. Adjusted p-values reflect BH FDR correction across the 53 networks.

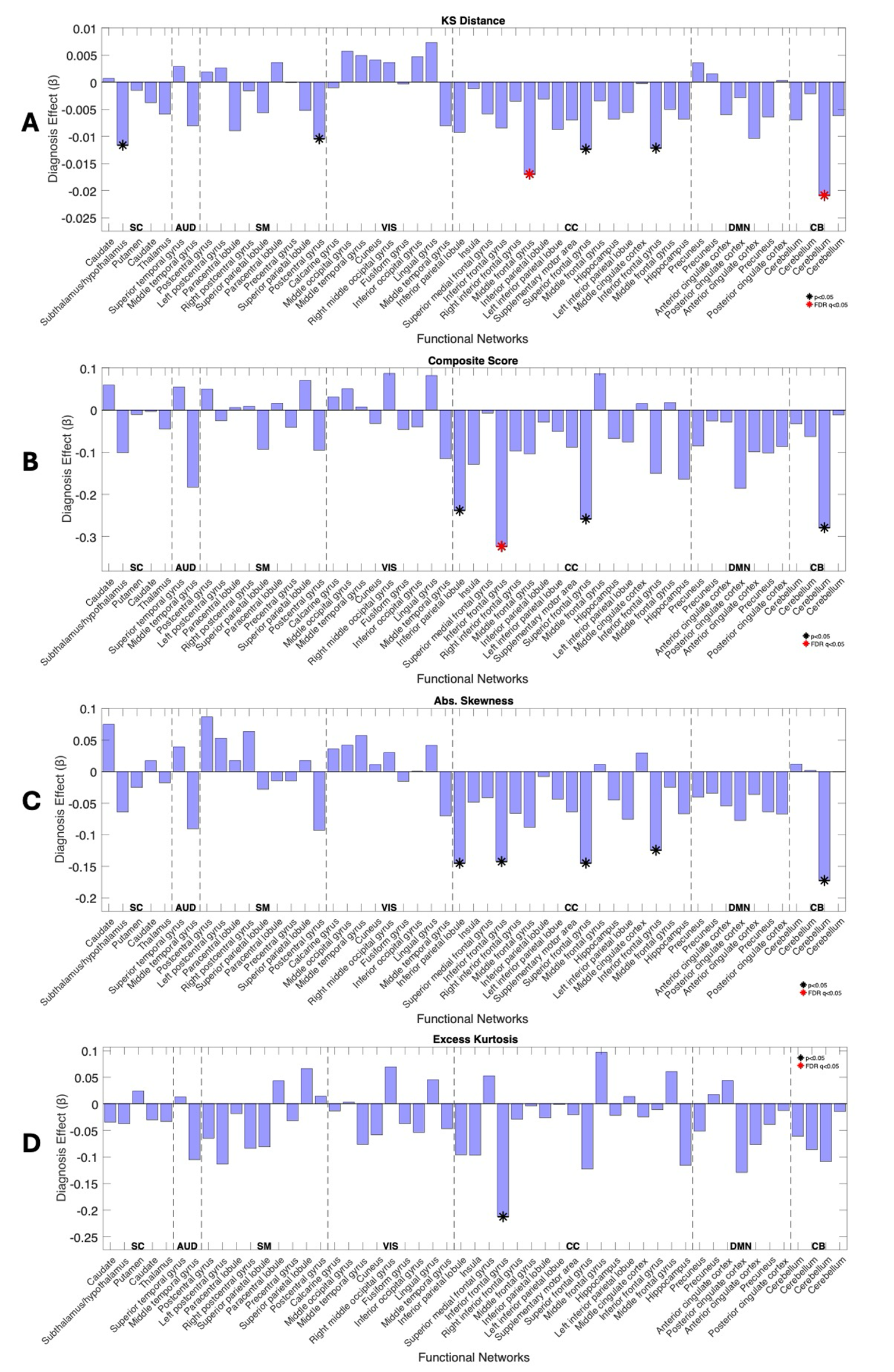

**Figure S3.**  Network-level diagnosis effect (β) on DICE distributions. Panels display β estimates for (A) KS distance, (B) Composite score, (C) Absolute skewness, (D) Excess Kurtosis. Black asterisks indicate p-values less than 0.05, red asterisks indicate p-values survived FDR correction; β<0 indicates lower values in SZ. Adjusted group differences were assessed with regression controlling for age, sex, and head motion. Adjusted p-values reflect BH FDR correction across the 53 networks.

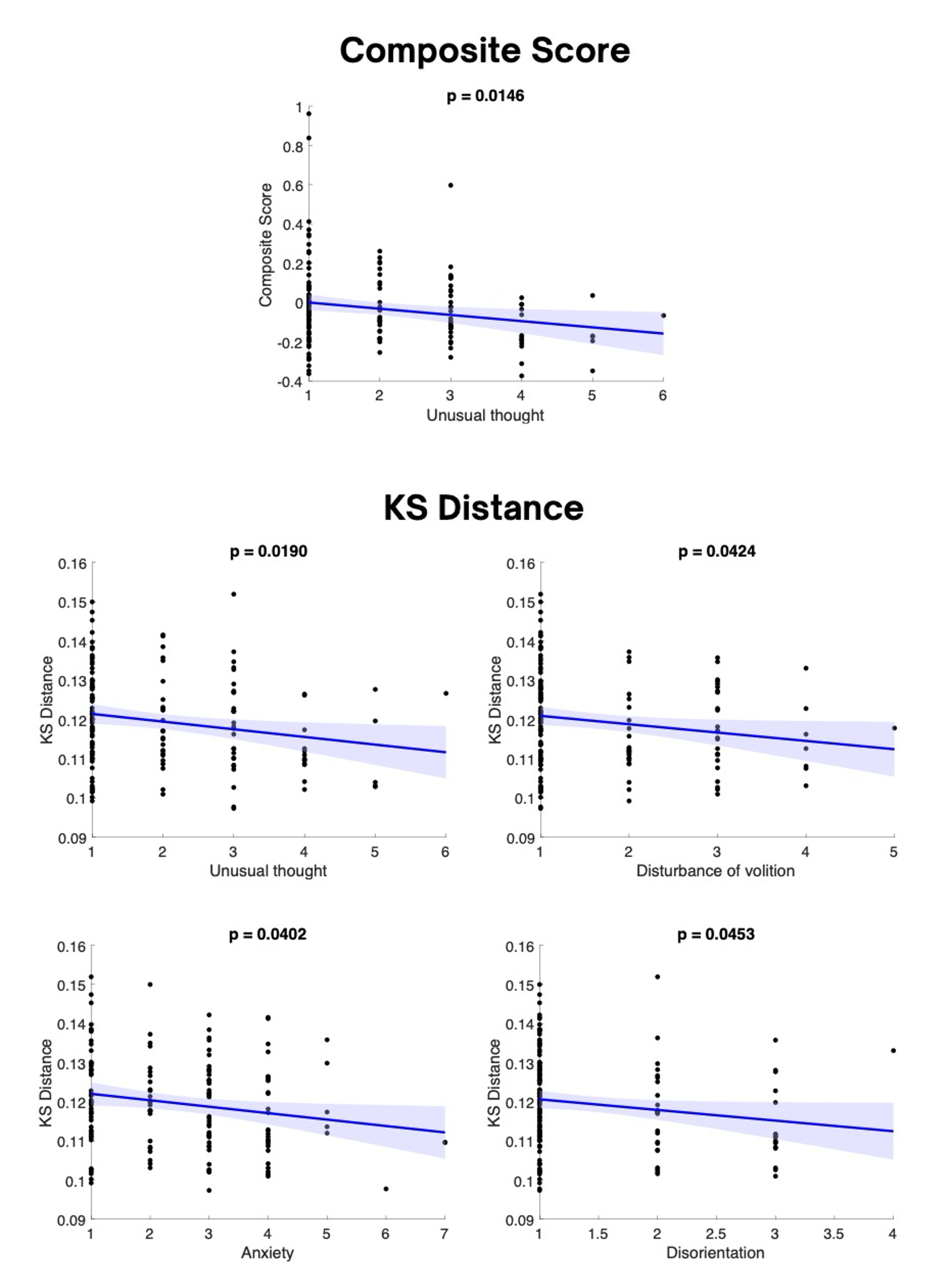

**Figure S4. (Supplementary to Figure 4)** Convergent distributional metrics (composite score and KS distance) at global level are associated with PANSS symptom severity in SZ. Solid lines show fitted linear regression trends; shaded bands indicate 95% confidence intervals. Associations were tested using linear regression controlling for age, sex, and head motion. Adjusted p-values reflect BH FDR correction across the 53 networks.

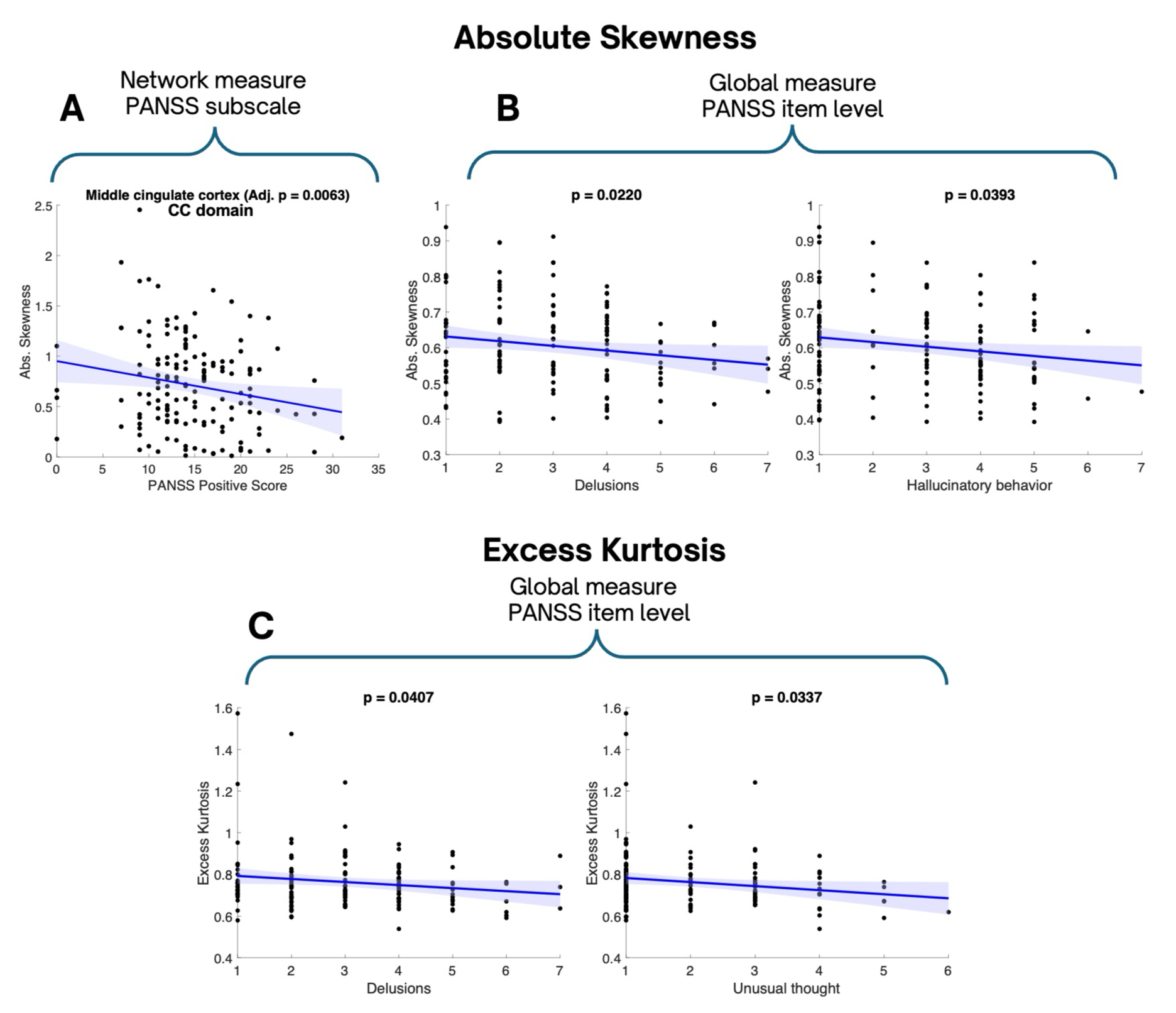

**Figure S5.** PANSS associations with non-Gaussianity metrics of dynamic entropy distributions (absolute skewness and excess kurtosis). (A) Network-level absolute skewness in the MCC (CC domain) is negatively associated with PANSS Positive subscale score. (B) Global (network-averaged) absolute skewness shows negative associations with the PANSS items delusions and hallucinatory behavior. (C) Global excess kurtosis shows negative associations with the PANSS items delusions and unusual thought content. Solid lines show fitted linear regression trends; shaded bands indicate 95% confidence intervals. Associations were tested using linear regression controlling for age, sex, and head motion. Adjusted p-values reflect BH FDR correction across the 53 networks.

.

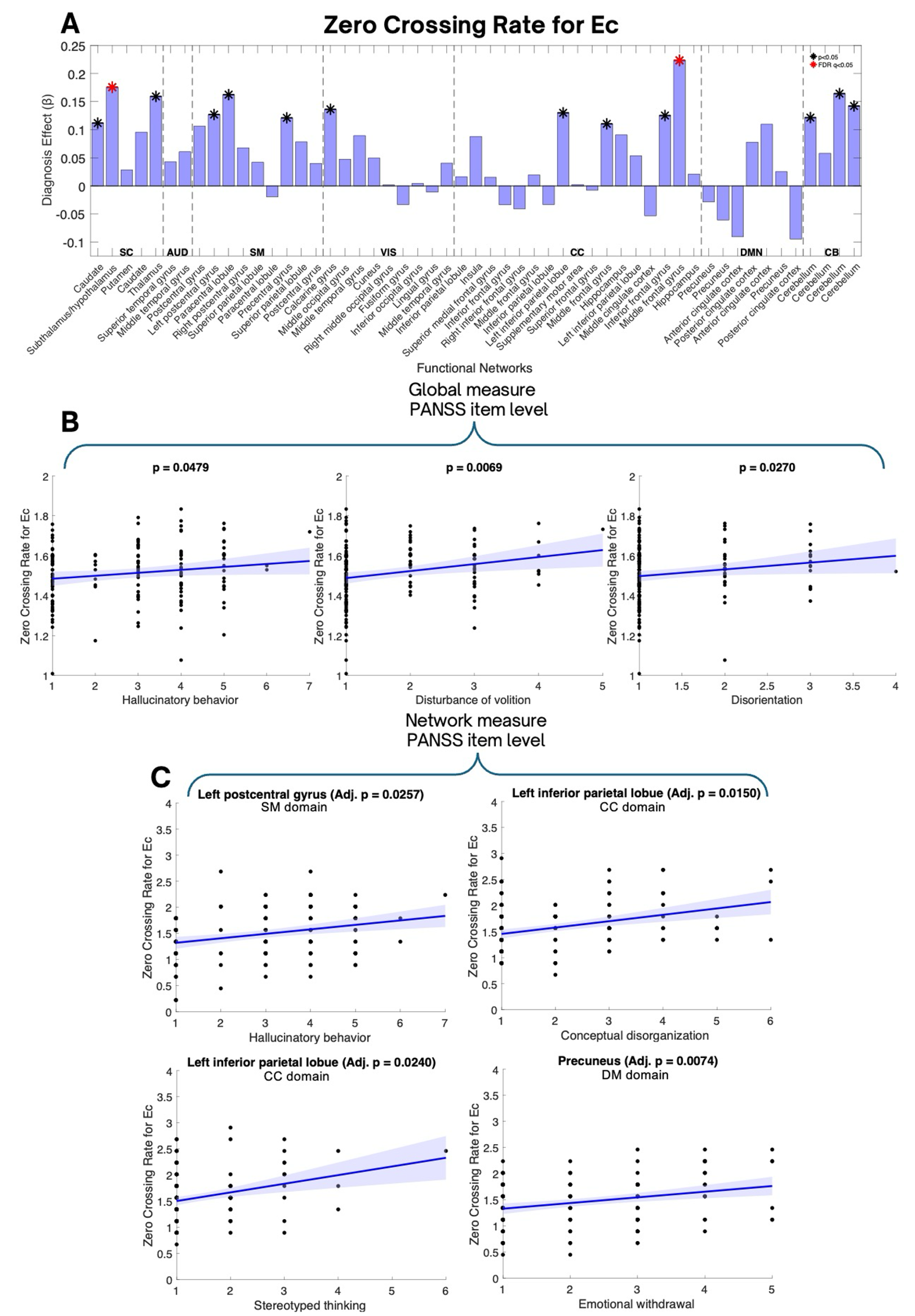

**Figure S6.** PANSS item-level associations with zero crossing rate of DICE fluctuations. Scatterplots show covariate-adjusted associations between PANSS item scores and the zero crossing rate of centered DICE fluctuations in SZ, with fitted regression lines and 95% confidence intervals. (A) Diagnosis effect, β, for mean zero-crossing rate; β>0 indicates higher values in SZ. Black asterisks show p-values less than 0.05, red asterisks indicate p-values survived FDR correction; (B) Global (network-averaged) zero crossing rate is positively associated with hallucinatory behavior, disturbance of volition, and disorientation. (C) Network-level zero crossing rate shows positive associations with symptom severity in selected regions; left postcentral gyrus (SM domain) with hallucinatory behavior, left inferior parietal lobule (cognitive control domain) with conceptual disorganization and stereotyped thinking, and precuneus (default mode domain) with emotional withdrawal. P-values correspond to the symptom term in regression models controlling for age, sex, and head motion; Adjusted p-values reflect BH FDR correction across the 53 networks.

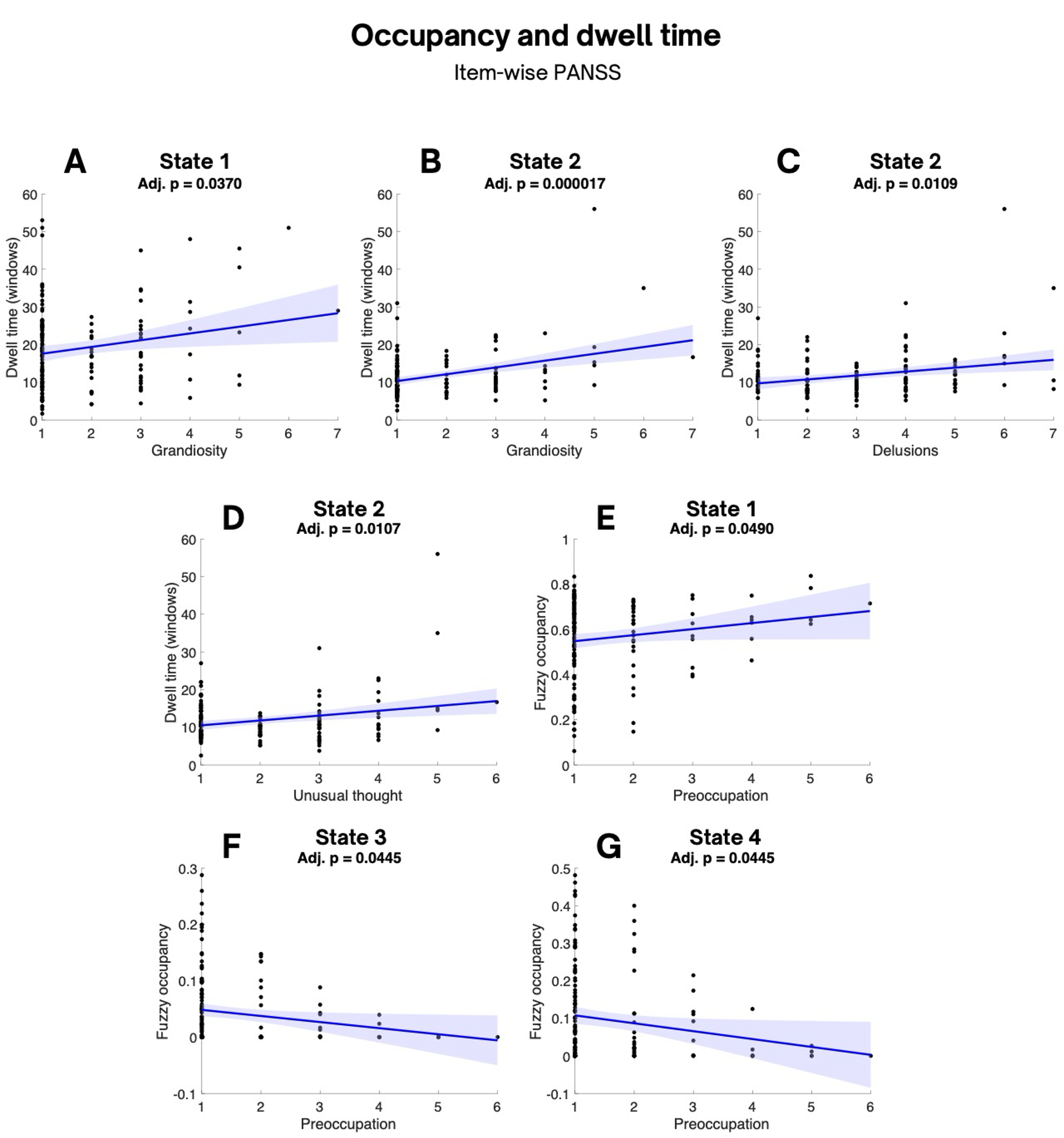

**Figure S7.** Deviation in entropy dwell time (A-D) and fuzzy occupancy (E-G) are linked to severity of diverse SZ symptoms in according with PANSS subscales. Symptoms associations were evaluated with linear regression approach controlling for age, sex, and head motion; FDR-adjusted p < 0.05 (BH). Solid lines indicate fitted regression trends with shaded 95% confidence intervals. Adjusted p-values reflect BH FDR correction across the four entropy states.
